# *Cyfip1* haploinsufficiency increases compulsive-like behavior and modulates palatable food intake: Implications for Prader-Willi Syndrome

**DOI:** 10.1101/264846

**Authors:** Richard K. Babbs, Qiu T. Ruan, Julia C. Kelliher, Jacob A. Beierle, Melanie M. Chen, Ashley X. Feng, Stacey L. Kirkpatrick, Fabiola A. Benitez, Fred A. Rodriguez, Johanne J. Pierre, Jeya Anandakumar, Vivek Kumar, Megan K. Mulligan, Camron D. Bryant

## Abstract

Binge eating (BE) is a heritable trait associated with eating disorders and involves rapid consumption of large quantities of food. We identified cytoplasmic FMRP-interacting protein 2 (*Cyfip2*) as a major genetic factor underlying BE and concomitant compulsive-like behaviors in mice. *CYFIP2* is a gene homolog of *CYFIP1 -* one of four paternally-deleted genes in patients with the more severe Type I Prader-Willi Syndrome (PWS). PWS is a neurodevelopmental disorder where 70% of cases involve paternal deletion of 15q11-q13. PWS symptoms include hyperphagia, obesity (if untreated), cognitive deficits, and obsessive-compulsive behaviors. We tested whether *Cyfip1* haploinsufficiency (+/-) would enhance premorbid compulsive-like behavior and palatable food (PF) intake in a parent-of-origin-selective manner. We tested *Cyfip1*^+/-^ mice on a C57BL/6N (N) background that were homozygous for the BE-associated missense mutation in *Cyfip2* (S968F) as well as mice that we backcrossed to homozygosity for the C57BL/6J (J) allele at *Cyfip2* (*Cyfip2*^J/J^). *Cyfip1*^+/-^ mice showed increased compulsive-like behavior on both backgrounds, increased PF consumption on the *Cyfip2*^N/N^ background in a paternally-enhanced manner, and decreased PF consumption in male *Cyfip1*^*+/-*^ mice on the *Cyfip2*^J/J^ background in a maternally selective manner. In the hypothalamus, there was a maternally-enhanced reduction of *Cyfip1* transcription, but a paternally-enhanced reduction in CYFIP1 protein. In the nucleus accumbens, there was a maternally-enhanced reduction in CYFIP1 protein. Together, increased compulsive-like behavior, parent-of-origin-, and genetic background-dependent effects of *Cyfip1* haploinsufficiency on PF consumption implicate *CYFIP1* in behaviors in neurodevelopmental disorders involving reduced expression of *CYFIP1*, including PWS, Fragile X Syndrome, and 15q11.2 Microdeletion Syndrome.

## INTRODUCTION

Binge eating (**BE**) refers to the rapid consumption of large quantities of food and is accompanied by feelings of loss of control. Binge eating disorder (**BED**) is a psychiatric disorder with a lifetime prevalence of 3.5% in women and 2% in men^1^. Both BED^2^ and BE are heritable^3^. However, genome-wide association studies have yet to identify genetic risk factors associated with BE^4^. The first genome-wide significant loci were recently identified for anorexia nervosa (comprising restricted eating)^5^ and bipolar disorder with BE behavior PRR5-ARHGAP8^6^. Additional genome-wide significant loci will likely be uncovered for BE-associated disorders with increasing sample sizes and power^7^.

We used quantitative trait locus (**QTL**) mapping and gene knockout in C57BL/6 mouse substrains to identify cytoplasmic FMR1-interacting protein 2 (*Cyfip2*) as a major genetic factor underlying BE and compulsive-like behaviors^8^. The QTL capturing increased palatable food (PF) intake mapped to a missense mutation in *Cyfip2* in the C57BL/6N strain (S968F; “*Cyfip2*^M1N^”) that is hypothesized to act as a gain-of-function mutation^9^. Accordingly, mice with one copy of a null allele and one copy of the missense allele of *Cyfip2* showed a reduction in BE toward the phenotypic direction of the wild-type C57BL/6J level^8^. This same missense SNP in *Cyfip2* was first associated with reduced behavioral sensitivity to cocaine^9^, which could indicate a common neurobiological mechanism involving synaptic plasticity within the mesocorticolimbic dopamine reward pathway^10,11^ that affects the hedonic component of PF consumption^12,13^.

*Cyfip2* and the gene homolog *Cyfip1* code for proteins that interact with the RNA binding protein Fragile X Mental Retardation Protein (**FMRP**) and are part of the canonical WAVE regulatory complex and transduce activity-dependent Rac signaling in regulating actin dynamics during neuronal development and synaptic plasticity^14^. CYFIP1 expression is necessary for the maintenance and stabilization of neuronal dendritic arborization and morphological complexity^15^. In humans, *CYFIP1* resides within a non-imprinted region on chromosome 15 (15q11.2) that contains four genes *TUBGCP5*, *NIPA1*, *NIPA2*, and *CYFIP1*^16^. The syntenic region in mice is located on chromosome 7C (55.4 Mb - 56 Mb). Haploinsufficiency of 15q11.2 underlies Microdeletion Syndrome (**MDS**) which can comprise developmental delay (speech, motor), reduced cognitive function, dysmorphic features, intellectual disability, autism, ADHD, obsessive-compulsive disorder, and schizophrenia^17^. At least one case study of 15q11.2 MDS reported hypotonia, increased food craving and obesity, and obsessive-compulsive disorder^18^. *CYFIP1* haploinsufficiency is implicated in multiple symptoms of 15q11.2 MDS. Preclinical models of *Cyfip1* haploinsufficiency demonstrate perturbations in synaptic activity during neural development, activity-dependent plasticity, dendritic morphology, and fear learning^19-22^.

The 15q11.2 region is also paternally-deleted in a subset of individuals with a more severe form (Type I) of Prader-Willi Syndrome (**PWS**), a neurodevelopmental disorder defined genetically by paternal deletion of 15q11-q13 in a majority of cases^23^. Extreme hyperphagia due to lack of satiety is the most defining and debilitating feature of PWS that is difficult to treat and emerges during childhood, leading to obesity if left untreated. Food-related obsessive-compulsive (**OC**) behaviors are common in PWS; however, OC symptoms unrelated to food are also frequent^24^, and include repetitive, ritualistic behaviors, perseverative speech, counting, adaptive impairment, need to tell, ask, or know, ordering and arranging, repeating rituals, and self-mutilation^25-27^. Genetic deletion in PWS involves either the shorter paternal deletion (Type II) of 15q11-q13 or a larger, paternal Type I deletion that also includes the 0.5 Mb 15q11.2 MDS region comprising four genes: *TUBGCP5*, *NIPA1*, *NIPA2*, and *CYFIP1*^16,28^. Type I PWS is associated with reduced transcription of these genes and a more severe neurodevelopmental and neuropsychiatric profile, including reduced cognition, increased risk of autism and schizophrenia, and increased severity and lack of control over OC behaviors (e.g., grooming and bathing, arranging objects, object hoarding, checking) that interfere with social functioning^16,18,28-30^.

Decreased CYFIP1 expression is also implicated in the Prader-Willi Phenotype (**PWP**) of a subset of individuals with Fragile-X Syndrome (**FXS**). FXS is the most common genetic cause of intellectual disability and autism and is caused by a CGG trinucleotide repeat expansion within the fragile X mental retardation 1 (*FMR1*) gene that is located on the X chromosome and codes for FMRP, a major interacting protein of CYFIP proteins^31^. Interestingly, ten percent of FXS individuals also exhibit a PWP in the absence structural or imprinting differences in 15q11-q13. The PWP includes hallmark hyperphagia, lack of satiation, obesity, and more severe behavioral problems, such as OC behaviors and an increased rate of autism^32,33^. The cause of the PWP is unknown, although one logical candidate gene is *CYFIP1*, given its association with PWS and its interaction with FMRP^31^. PWP-presenting individuals with FXS show a two-to four-fold decrease in CYFIP1 transcription compared to FXS individuals without PWP^33^. There was also a two-fold decrease in *Cyfip1* gene transcription in a mouse model of FXS^34^.

Because of the association of the gene homolog *Cyfip2* with BE^8^ and because both *CYFIP1* deletion and reduced CYFIP1 expression are associated with PWS (Type I) and hyperphagia in the PWP (FXS) respectively, here, we tested the hypothesis that *Cyfip1* haploinsufficiency would increase premorbid compulsive-like behavior and consumption of palatable food (**PF**) in our BE paradigm^8,35,36^. We tested the effect of *Cyfip1* deletion on two different *Cyfip2* genetic backgrounds. Additionally, because a recent study demonstrated a parental origin (**PO**) effect of *Cyfip1* haploinsufficiency on hippocampal synaptic transmission, learning, and anxiety-like behavior^19^, we tested for a PO effect of *Cyfip1* deletion on compulsive-like behavior and PF intake. To gain insight into the molecular mechanisms underlying the PO effect of *Cyfip1* deletion on PF intake, we examined transcription of *Cyfip1*, *Cyfip2*, and *Magel2* - a nearby imprinted gene within the syntenic, canonical PWS region implicated in hyperphagia and obesity. Additionally, we examined CYFIP1 protein expression in the hypothalamus and nucleus accumbens as a function of both *Cyfip1* genotype and PO. Finally, because OC behaviors are associated with BE^37-39^ and hyperphagia in PWS^40^, we employed a battery of tests to assess anxiety-like and compulsive-like behaviors and post-BE training behaviors, including compulsive-like eating and concomitant behaviors in the light/dark conflict test^8^ in *Cyfip1* haploinsufficient mice.

## MATERIALS AND METHODS

### Mice

All experiments were performed in accordance with the National Institutes of Health Guidelines for the Use of Laboratory Animals and were approved by the Institutional Animal Care and Use Committee at Boston University. Mice were 50-100 days old at the first day of testing. A minimum sample size of N = 20 per Genotype per Treatment was employed for behavioral studies based on power analysis of PF intake from the *Cyfip2* study^8^ (see **Supplementary Information** for additional details). Mice heterozygous for a null deletion in exons 4 through 6 of *Cyfip1* (*Cyfip1*^+/-^) were propagated on two different C57BL/6 genetic backgrounds (**Fig.1)**: (1) the BE-prone C57BL6/N background, and (2) a mixed background whereby mice were homozygous for the BE-resistant C57BL6/J background at the *Cyfip2* locus. Details regarding generation of mice and *Cyfip1* and *Cyfip2* genotyping are provided in the **Supplementary Information**.

### Premorbid anxiety- and compulsive-like behavioral battery

Because of the link between anxiety, compulsivity and pathological overeating^39^ and because OC behavior is associated with eating disorders^41,42^, we incorporated a behavioral battery to assess differences in premorbid anxiety-like and compulsive-like behaviors in experimentally naïve, *Cyfip1*^+/-^ mice. Mice were tested in the behavioral battery and were either sacrificed afterward (mice on *Cyfip2*^N/N^ background) or were subsequently trained for BE (mice on the *Cyfip2*^J/J^ background). Mice were assayed in the battery with one test per day over five days in the following order: 1) open field; 2) elevated plus maze; 3) marble burying; 4) hole board; 5) mist-induced grooming. Procedural details are provided in the **Supplementary Information**. Testing was conducted between 0800 and 1300 h. The experimenters responsible for running the mice, video tracking, data curation, and analysis were blinded to Genotype for each cohort.

Digging, burrowing, and burying of objects with bedding are highly correlated behaviors associated with survival. Inherent burying of non-aversive stimuli can be distinguished from defensive burying of aversive, noxious stimuli^43^ and is not related to anxiety, stimulus novelty, or locomotor activity levels. Rather, inherent marble burying is an indirect measure of the natural tendency to dig^44-46^. Marble burying represents persistent, repetitive behavior that is resistant to habituation and is proposed to model obsessive/compulsive behavior^45,47^. This behavior is used to screen pharmacotherapeutics for OCD^48^. Genetic variance in marble burying among inbred mouse strains is heritable but is genetically uncorrelated with anxiety-like behaviors and is thus, mediated by distinct genetic factors^45^.

The hole board test is commonly used to assess anxiety^49^, novelty-seeking^50^, and repetitive behavior^51^. Moreover, head-dipping activity in the hole board test has been shown to be a valid predictor of reward-associated behaviors such as nicotine self-administration^52^, and cocaine-induced conditioned place preference^50^.

### BE procedure and light/dark conflict test of compulsive eating

Mice were trained in an intermittent, limited access procedure to detect genetic differences in BE^8,35^. For details, see **Supplementary Information**. Briefly, mice were tested for side preference on D1 and D22. In the intervening days, mice were confined to a food-paired and non-food-paired side on alternating days (Tues-Fri). Cages were assigned to either the PF or Chow group in a counterbalanced design in order to ensure equal distribution across Sex, Genotype, Treatment, and PO. On D23, mice were assessed for compulsive eating and associated behaviors, as previously described^8^ (**Supplementary information**). The experimenters responsible for running the mice, video tracking, data curation, and analysis were blinded to Genotype for each cohort.

### Hypothalamus dissections for real-time quantitative PCR (qPCR)

We chose a subset of Chow-trained, PF-naive mice (n = 7-9 per Genotype per PO; both sexes) on the *Cyfip2*^N/N^ background or untrained, PF-naïve mice on a *Cyfip2*^J/J^ background (n= 8-12 per Genotype per PO; both sexes) to examine baseline (PF-naive) gene transcription between *Cyfip1*^N/-^ versus *Cyfip1*^N/N^ mice and PO. We examined *Cyfip1*, *Cyfip2*, and *Magel2* transcript levels in the hypothalamus, a brain region important for hyperphagia in PWS ^40^ and for the effects of *Magel2* deletion^53,54^ on eating behavior and homeostatic function^55^. Haploinsufficiency of *MAGEL2* is associated with PWS-like hyperphagia in humans^54,56,57^.

On D24, brains from Chow-trained mice (*Cyfip2*^N/N^ background) were harvested and the hypothalamus was free form dissected by pinching the entire structure from the ventral surface with forceps while using the anterior commissure and mammillary bodies as landmarks. Tissue was stored in RNAlater Solution (Invitrogen, Carlsbad, CA USA) at 4°C. After five days, the tissue was dried and transferred to a −80°C freezer.

### Real-time quantitative PCR (qPCR)

Total RNA from hypothalamus was extracted and processed for qPCR as described^8,35,58^. Briefly, oligo-dT primers were used to synthesize cDNA. PCR reactions were conducted on the StepOne Plus 96-Well Real-Time PCR machine (Life Technologies, Foster City, CA, USA) in technical triplicates and averaged (SD < 0.5). Plates were balanced across Genotype, PO, and Sex. We report the difference in expression in *Cyfip1*^+/-^ relative to *Cyfip1*^+/+^ using the 2^−(ΔΔCT)^ method^59^. Primer sequences are provided in the **Supplementary Information**.

### Western blot of hypothalamus and nucleus accumbens

Hypothalamus was dissected as described above. Nucleus accumbens punches were harvested using 1.2 mm punches of ventral forebrain centered around anterior commissure from the first 4 mm of brain section in a brain matrix. Samples were processed and analyzed for quantity of CYFIP1 protein. Detailed methods can be found in the **supplementary material**. Because we found no effect of Treatment for either brain region, data were collapsed across Treatment for analysis.

### Analysis

Statistical analyses were conducted using R (https://www.r.project.org). For the compulsive behavioral tests, two-tailed unpaired t-tests were used to detect effects of Genotype for all behaviors except marble burying behaviors which were analyzed by non-parametric Mann-Whitney U tests. Slope analyses were conducted as previously described^8,60^ using GraphPad Prism 7 (GraphPad Software, La Jolla, CA USA). We analyzed food intake using mixed model ANOVAs with Genotype, Treatment, and Sex as independent variables, and Day as a repeated measure using the “aov” function in R. Additionally, we assessed the effect of PO (maternal, paternal) using mixed-model ANOVAs with Genotype, Treatment, and PO as independent variables and Day as a repeated measure. To address issues of non-normality or unequal variance, we included additional non-parametric analyses to support key findings (**Supplementary Results**).

## RESULTS

### *Cyfip1* haploinsufficiency increases compulsive-like behaviors

Sample sizes are listed in **Supplementary Table 1**. **Figure 1** illustrates the breeding scheme for *Cyfip1*^+/-^ mice on two *Cyfip2* genetic backgrounds: *Cyfip2*^N/N^ and *Cyfip2*^J/J^. Because symptomatic severity is worse in individuals with Type I PWS (which includes the *CYFIP1* deletion) and because compulsivity was negatively correlated with *CYFIP1* expression in PWS patients with Type I deletions^16^, we hypothesized that *Cyfip1*^+/-^ mice would exhibit greater compulsive-like behavior.

**Figure 1.**
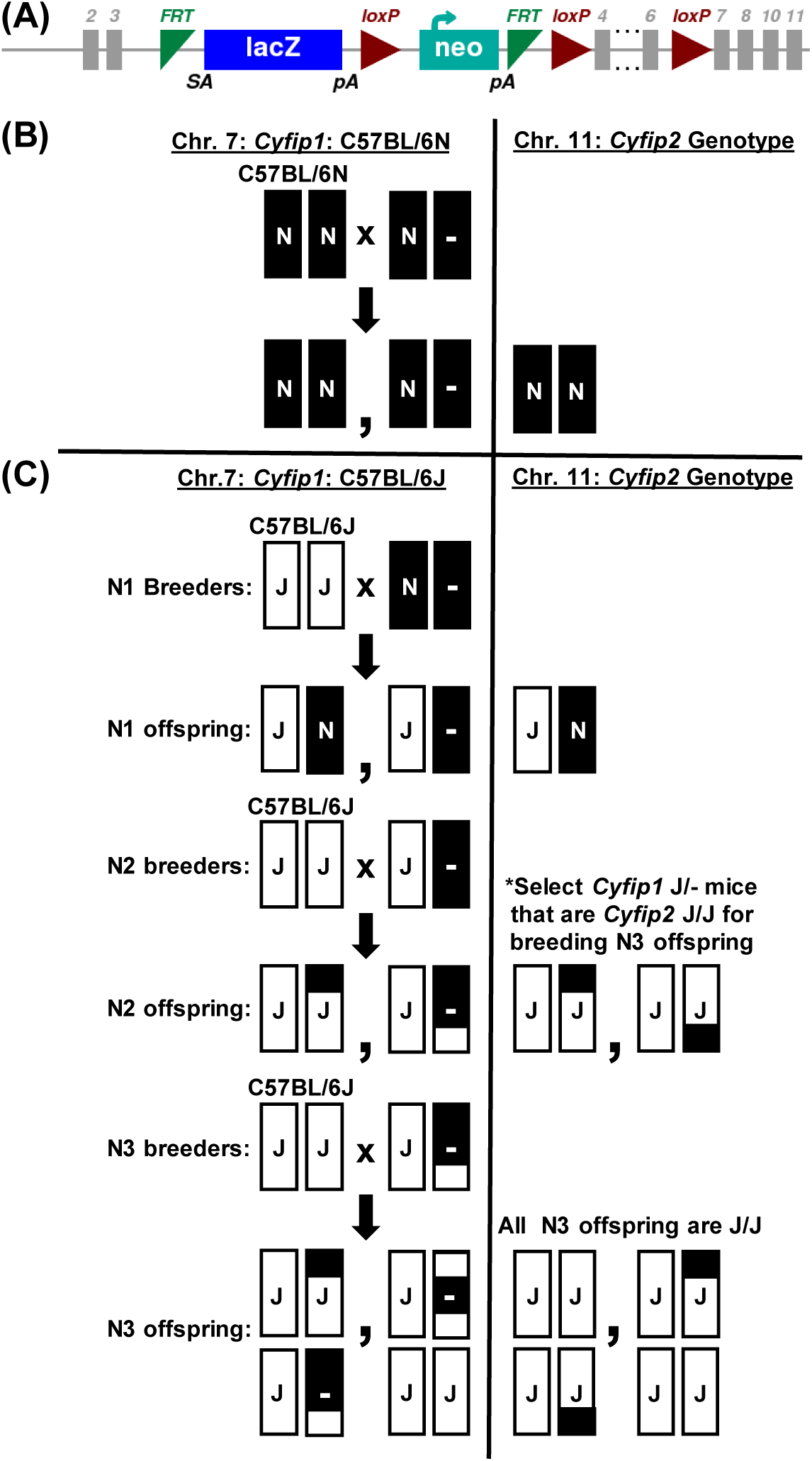
Generation of the *Cyfip1* knockout allele and breeding scheme for *Cyfip1* haploinsufficient mice on *Cyfip2*^N/N^ and *Cyfip2*^J/J^ genetic backgrounds. **(A):** A schematic of the knockout first allele for KOMP generation of *Cyfip1*^N/-^ mice was obtained from the International Mouse Phenotyping Consortium (IMPC) website (http://www.mousephenotype.org/data/alleles/MGI:1338801/tm2a(EUCOMM)Wtsi). Mice containing floxed alleles flanking exons 4 through 6 were generated from embryonic stem cells on a C57BL/6N background by the International Knockout Mouse Consortium and were crossed to global Cre-expressing mice, yielding offspring heterozygous for constitutive deletions in exons 4 through 6. We derived mice heterozygous for the null deletion on a C57BL/6N background using sperm obtained from The Jackson Laboratory. **(B): Left panel:** In the first study, we re-derived *Cyfip1*^N/-^ mice on an isogenic C57BL/6N background. **Right panel:** All mice were homozygous for the N allele (N/N) at *Cyfip2* which contains a missense mutation that we previously showed was associated with a marked enhancement of binge eating (BE), accounting for approximately 50% of the genetic variance in parental strain BE^8^. We maintained this colony on an isogenic C57BL/6N background by breeding *Cyfip1*^N/-^ mice with C57BL/6NJ mice (black bars; N/N) ordered from The Jackson laboratory. **(C):** In the second study, we generated another colony on a mixed background. The primary goal was to monitor and replace the BE-associated N/N *Cyfip2* alleles with C57BL/6J (J/J) alleles via backcrossing *Cyfip1*^N/-^ mice to C57BL/6J (white bars; J/J) for three and four generations and assess the effect of *Cyfip1* deletion on BE on a mixed N3 and N4 background containing a fixed, BE-resistant, homozygous J/J genotype at *Cyfip2* ^8^. Mixed-color bars illustrate hypothetical recombination events that accumulate through backcrossing to C57BL/6J (white).

In the marble burying test, *Cyfip1*^N/-^ mice on the *Cyfip2*^N/N^ genetic background showed a greater number of marbles that were at least 50% buried and a greater average percentage of marbles buried than *Cyfip1*^N/N^ mice (**Fig.2A-C**) which are commonly used measures of marble burying^61^. This result was replicated in *Cyfip1*^J/-^ on the *Cyfip2*^J/J^ background (**Fig.2D,E**). *Cyfip1* deletion on the *Cyfip2*^N/N^ genetic background did not induce a change in any other behaviors within the battery (**Supplementary Table 2**, all ps > 0.05). However, on the *Cyfip2*^J/J^ background, *Cyfip1*^J/-^ showed a greater number of head dips in the hole board test than *Cyfip1*^J/J^, (**Supplementary Fig.1C**), further supporting increased compulsive-like behaviors induced by *Cyfip1* haploinsufficiency.

**Figure 2.**
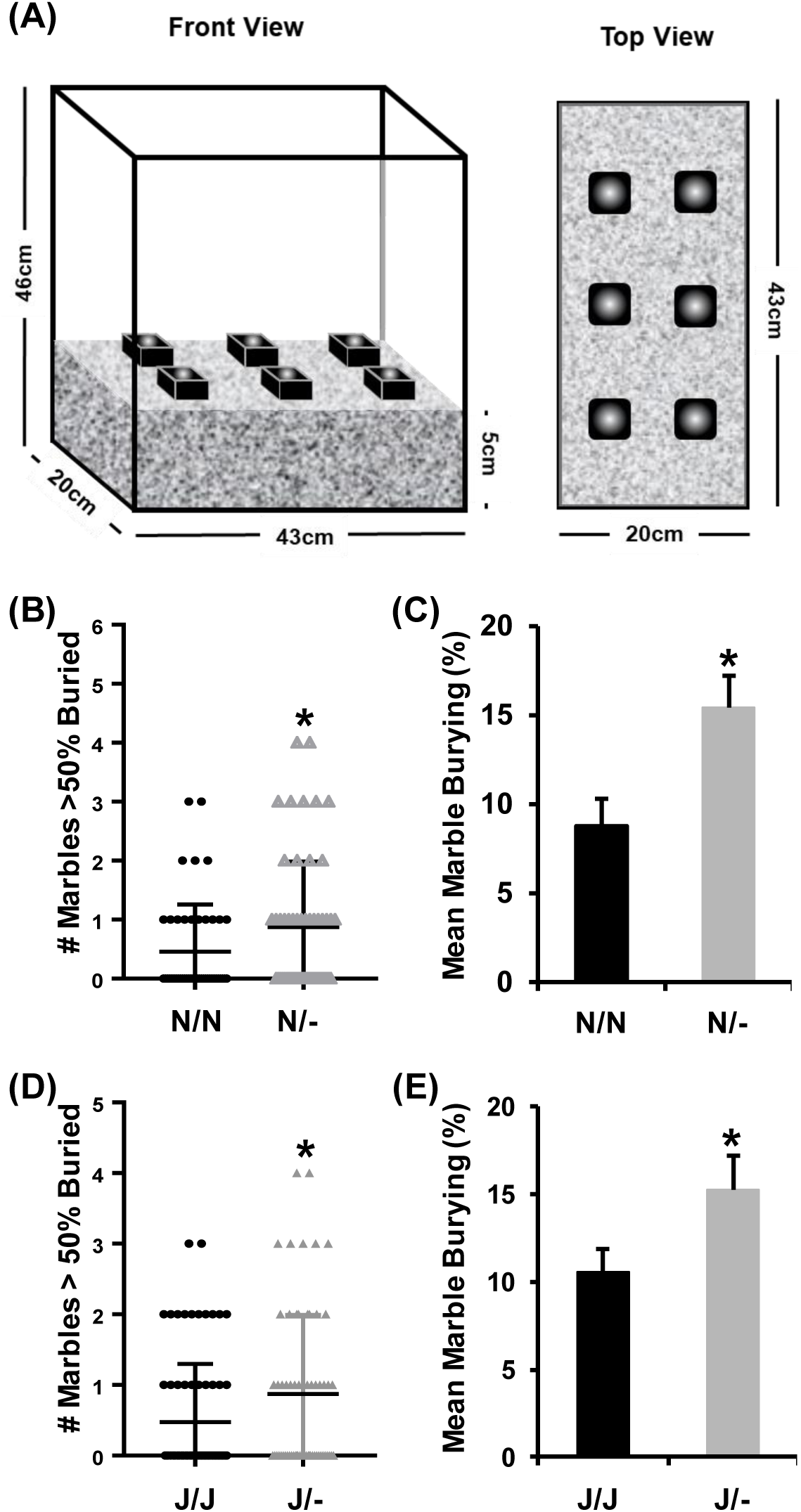
Increase in premorbid, OC-like marble burying in *Cyfip1*^N/-^ and *Cyfip1*^J/-^ mice. **(A):** Schematic of the marble burying apparatus. **(B,C):** *Cyfip1*^N/-^ mice buried more marbles with greater than 50% coverage than wild-type *Cyfip1*^N/N^ mice [B: U(102) = 1050; p = 0.031; two-tailed], and had a greater average percentage of burial across the marbles [**C**; F(1,96) = 7.1; p = 0.009]. **(D,E):** Similarly, *Cyfip1*^J/-^ mice buried more marbles with greater than 50% coverage than J/J mice [**D:** U(137) = 1884; p = 0.019, two-tailed], and also had a greater average percentage of burial [**E**; F(1,134) = 4.2; p = 0.042]. Data are presented as mean ± SEM.

Because *CYFIP1* is paternally-deleted in Type I PWS and because PO effects of *Cyfip1* deletion on synaptic transmission and behavior were reported^19^, we next investigated the effect of PO of *Cyfip1* deletion on anxiety-like and compulsive-like behaviors. There was no effect of PO or interaction with *Cyfip1* Genotype on marble burying or any other behaviors within the battery (data not shown). To summarize, *Cyfip1* haploinsufficiency induced a selective increase in compulsive-like marble burying regardless of genetic background, as well as an increase in compulsive-like head-dipping in the hole board test in mice on the *Cyfip2*^J/J^ genetic background.

### Effect of *Cyfip1* haploinsufficiency on PF intake depends on *Cyfip2* genetic background

In testing the hypothesis that *Cyfip1* haploinsufficiency would increase PF intake in our intermittent, limited access BE and CPP paradigm (**Fig.3A**), we first found that PF-trained mice of both genotypes on the *Cyfip2*^*N/N*^ background consumed significantly more food than Chow-trained mice (**Fig.3B**) – this result was reflected by slopes of escalation that were significantly greater than zero in both PF-trained genotypes but not in the Chow-trained genotypes (**Fig.3C**). As predicted, *Cyfip1*^N/-^ mice consumed more PF than *Cyfip1*^N/N^ mice, but not more Chow (**Fig.3B**). Finally, PF-trained *Cyfip1*^N/-^ mice showed a greater y-intercept than all other groups (**Fig. 3C**), indicating an initial higher level of consumption that persisted throughout the study.

**Figure 3.**
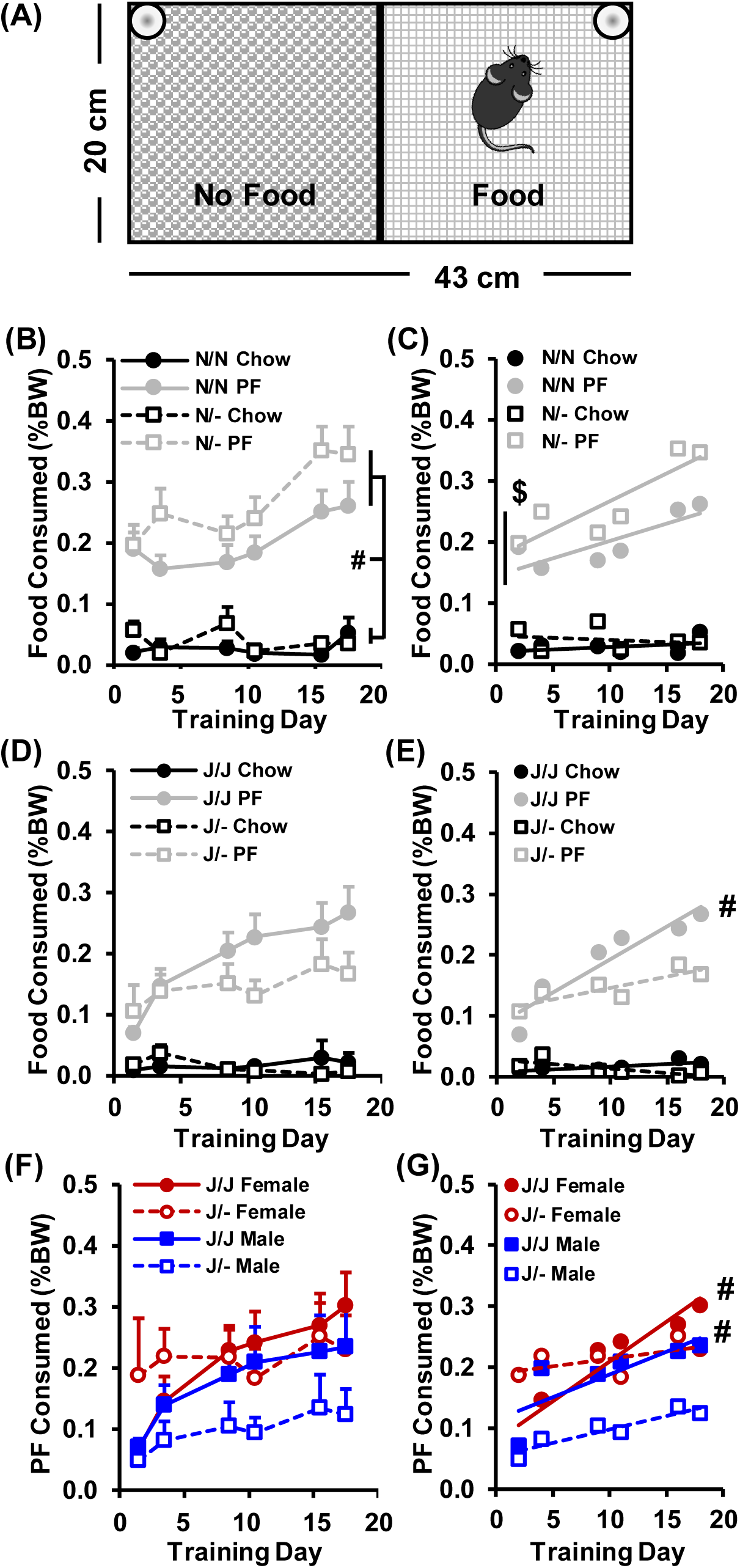
PF consumption in *Cyfip1*^N/-^ and *Cyfip1*^J/-^ mice. **(A:)** The conditioned place preference chamber used for food consumption training had a smooth-textured non-food-paired side (left) and a rough-textured food-paired side. **(B):** Both wild-type *Cyfip1*^N/N^ and *Cyfip1*^N/-^ mice trained with PF in the CPP chamber ate more food over time than Chow-trained mice [**#** main effect of Treatment: F(1,932) = 274.7; p < 2×10^−16^]. There was also a main effect of Sex [F(1,932) = 30.4; p = 4.5 x 10^−8^], a Genotype x Treatment interaction [F(1,932) = 4.7; p = 0.03], and a Treatment x Sex interaction [F(1,932) = 22.3; p = 2.7 x 10^−6^]. **(C):** Both PF-trained genotypes exhibited slopes that were significantly greater than zero (*Cyfip1*^N/N^: m = 0.009 ± 0.003, p = 0.024; *Cyfip1*^N/-^: m = 0.005 ± 0.002, p = 0.045, respectively) indicating escalation in PF intake over time. Neither Chow-trained group showed escalation. Moreover, PF-trained *Cyfip1*^N/-^ mice showed a significantly greater y-intercept than all other groups (**$**: p < 0.008), indicating consistently greater overall food consumption throughout the study. **(D):** When we examined the same behaviors in *Cyfip1*^J/-^ versus *Cyfip1*^J/J^ mice, we observed a main effect of Treatment [F(1,824) = 200.4; p = 2 x 10^−16^], indicating that PF-trained mice consumed more food over time. We also observed a main effect of Genotype [F(1,824) = 4.2; p = 0.04], but in contrast to *Cyfip1* ^N/-^ mice, *Cyfip1*^*J/-*^ consumed *less* food than their wild-type *Cyfip1*^*J/J*^ counterparts. Additionally, we observed a main effect of Sex [F(1,542) = 27.0, p = 3.0 x 10^−7^], Day [F(1,824) = 17.9; p = 2.6 x 10^−5^], a Sex x Treatment interaction [F(1,824) = 6.3; p = 0.01], a Treatment x Day interaction [F(1,824) = 16.9; p = 4.3 x 10^−5^], a Genotype x Day interaction [F(1,824) = 6.7; p = 0.01], and most importantly, a Sex x Treatment x Genotype interaction [F(1,824) = 11.9; p = 0.0006]. **(E):** In examining escalation in food intake over time, only PF-trained *Cyfip1*^*J/J*^ mice exhibited a positive non-zero slope [**#**; F(1,244) = 21.0; p < 0.0001], indicating escalation in consumption over time and further supporting reduced food intake in *Cyfip1*^J/-^ mice. **(F,G):** In order to better understand the interactions with Sex in PF-trained mice, we examined PF consumption in *Cyfip1*^*J/-*^ versus *Cyfip1*^*J/J*^ mice in females and males. We found main effects of Sex [F(1,446) = 11.2; p = 0.0009], Genotype [F(1,446) = 4.2; p = 0.04], and Day [F(1,446) = 21.3; p = 5.2 x 10^−6^]. We also observed a Genotype x Day interaction [F(1,446) = 4.8; p = 0.03], a Sex x Genotype interaction [F(1,446) = 7.8; p =0.005], a Gene x PO interaction [F(1,446) = 10.6; p =0.001], and a Sex x Genotype x PO interaction [F(1,446) = 5.7; p = 0.02]. **(G)** In examining escalation in PF consumption, we found that both female and male wild-type *Cyfip1*^*J/J*^ mice had positive slopes (**#**; both ps < 0.009 vs. zero) whereas both female and male *Cyfip1*^J/-^ mice did not have any significant slope (both ps > 0.08). However, the *Cyfip1*^*J/-*^ females had the greatest y-intercept (p < 0.0001) indicating the greatest initial consumption. Data are presented as mean ± SEM.

In examining the effect of *Cyfip1* haploinsufficiency on food intake on a *Cyfip2*^J/J^ genetic background, we observed less overall PF intake, as expected, relative to the *Cyfip2*^N/N^ background [t(155) = 2.4; p = 0.02]^8^. Moreover, PF-trained mice again showed greater intake than Chow-trained mice (**Fig.3D**). However, when examining the effect of Genotype on the *Cyfip2*^J/J^ background, the results were unexpected. Interestingly, *Cyfip1*^J/-^ mice showed *less* PF intake than their wild-type *Cyfip1*^J/J^ counterparts (**Fig.3D**) and did not escalate over time whereas wild-type mice escalated as indicated by a significant, positive slope (**Fig.3E**).

To further dissect the unexpected decrease in PF intake in *Cyfip1*^J/-^ mice, we examined female and males separately. Overall, there was greater PF intake in females compared to males **(Fig.3F)**. Surprisingly, male *Cyfip1*^J/-^ mice completely accounted for the decrease in PF consumption in *Cyfip1*^J/-^ mice (**Fig.3F**). Furthermore, neither female nor male *Cyfip1*^J/-^ showed an escalation in PF consumption whereas their wild-type *Cyfip1*^J/J^ counterparts of both sexes showed positive slopes (**Fig.3G**).

### PO-and sex-dependent effects of *Cyfip1* haploinsufficiency on PF intake

We next investigated the effect of PO of *Cyfip1* deletion on food intake in light of a recent study demonstrating a PO effect of *Cyfip1* deletion on emotional learning and synaptic transmission^19^. We focused on PF intake rather than Chow intake based on the above results.

For mice on the *Cyfip2*^N/N^ background, paternal *Cyfip1* deletion induced greater PF consumption in all offspring (**Figs.4A,B**). Paternally-deleted *Cyfip1*^N/-^ mice showed a greater y-intercept than both *Cyfip1*^N/N^ groups (**Fig.4C**), indicating overall greater intake across time. Wild-type *Cyfip1*^N/N^ offspring derived from families with paternal *Cyfip1* deletion also showed a greater y-intercept (**Fig.4C**), thus confirming an overall effect of parental *Cyfip1* genotype on neurobehavioral expression of all offspring within those families.

In examining PO effects in *Cyfip2*^J/J^ mice, maternal *Cyfip1* deletion accounted for the overall Genotype effect of decreased PF intake (**Fig4.D,E**). Additionally, only wild-type mice showed evidence for escalated intake and *Cyfip1*^J/-^ mice showed no significant slope, regardless of PO. Because we identified a Sex x Genotype x PO x Day interaction in mice with the *Cyfip2*^J/J^ background, we sought to identify the source of these interactions. When separated by Sex, paternally-deleted female *Cyfip1*^J/-^ mice showed *enhanced* PF intake as indicated by an increase in y-intercept (**Fig.4G-I**) whereas maternally-deleted male *Cyfip1*^J/-^ mice showed markedly *decreased* PF intake as indicated both by reduced PF consumption and no escalation over time (**Fig.4J-L**). To summarize, we observed markedly different effects of *Cyfip1* deletion on PF intake that depended on *Cyfip2* genetic background, PO, and Sex. Despite changes in PF intake across Genotype and PO, we did not detect any effects of Genotype or PO on body weight. (**Supplementary Fig.2**). Thus, homeostatic differences are unlikely to explain the above findings.

### Conditioned food reward in *Cyfip1*^+/-^ mice

In examining CPP on the *Cyfip2*^N/N^ genetic background, there was no effect of *Cyfip1* Genotype or Treatment in the ANOVA model in mice from either genetic background (**Supplementary Fig.3A,B**). However, when we considered PF treatment alone, there was increased PF-CPP in *Cyfip1*^N/-^ versus *Cyfip1*^N/N^ mice that was in line with increased PF consumption (**Supplementary Fig.3A**). For *Cyfip1*^J/-^ mice on the *Cyfip2*^J/J^ background, there was no genotypic difference (**Supplementary Fig.3B**). In considering PO effects on PF-CPP, there was no effect of Genotype, PO, or interaction on either *Cyfip2* genetic background (**Supplementary Fig.3C,D**).

### Compulsive-like eating in the light/dark conflict test in *Cyfip1*^+/-^ mice

Because we observed increased compulsive-like behavior in *Cyfip1*^+/-^ mice and because increased PF consumption can become compulsive-like^39^, we next examined post-training compulsive-like eating using the light/dark conflict test (**Fig.5A**)^8^. PF-trained mice showed greater compulsive-like consumption than Chow-trained mice (**Fig.5B,D**). Furthermore, females showed greater PF consumption than males on both genetic backgrounds (**Fig.5C,E**). *Cyfip1* deletion on the *Cyfip2*^N/N^ background had no effect on PF consumption (**Fig.5B,C**), regardless of PO (**Supplementary Fig.4A-C**). For the *Cyfip2*^J/J^ background, *Cyfip*^J/-^ mice showed reduced PF consumption that was driven primarily by the males (**Fig.5D,E**), similar to PF intake during training (**Fig.3F,G**). However, unlike the PO dependency of the reduced PF observed during training (**Fig.4J-L**), the male-specific reduction in PF intake during the compulsive-like eating test did not depend on PO (**Supplementary Fig.4D,E).**

**Figure 4.**
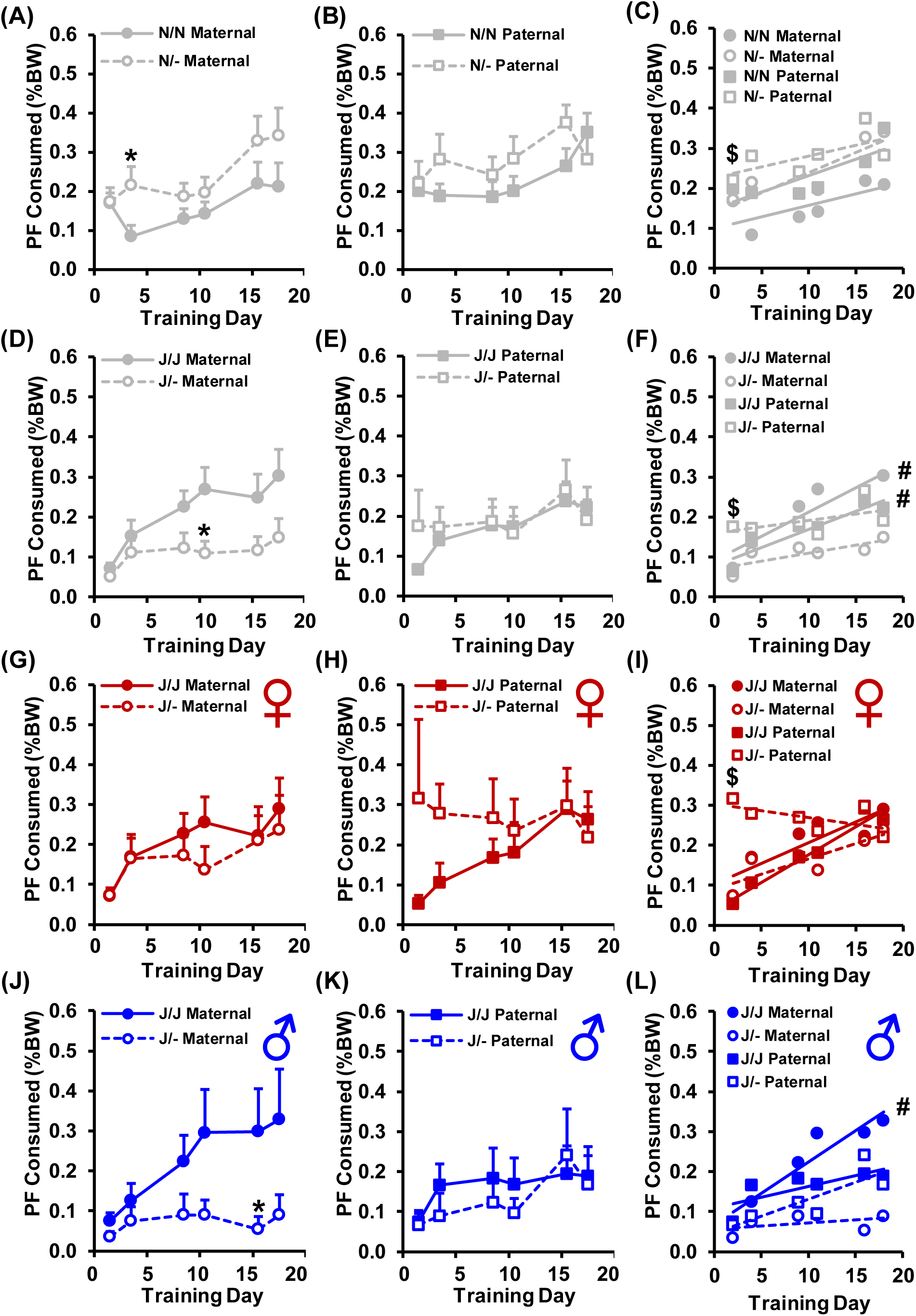
Effect of parent-of-origin (PO) on PF consumption in *Cyfip1*^N/-^ and *Cyfip1*^J/-^ mice. **(A,B):** In considering mice on a *Cyfip2*^N/N^ background, there was a main effect of Genotype [F(1,464) = 12.3; p = 0.0005], PO [F(1,464) = 9.0; p = 0.003], Sex [F(1,464) = 39.4; p = 8.0 x 10^−10^], and Day [F(1,464) = 20.8, p = 6.5 x 10^−6^]. The effect of Genotype was explained by *Cyfip1*^N/-^ mice consuming more PF than *Cyfip1*^N/N^ mice [*; t(29) = 2.1; uncorrected p = 0.046 vs. *Cyfip1*^N/N^ on D4]. The effect of Sex was explained by females consuming more PF than males (not shown). The effect of PO was explained by offspring derived from parents with a paternal *Cyfip1* deletion **(B)** consuming a greater amount of PF than offspring derived from parents with a maternal *Cyfip1* deletion **(A)**. **(C):** No differences were observed among the groups in the slopes of escalation in PF consumption [F(3,16) = 0.7 p = 0.56]; however, paternally-deleted *Cyfip1*^N/-^ mice (open squares) showed a greater overall consumption than either of the *Cyfip1*^N/N^ wild-type groups as indicated by a greater y-intercept (**$**: both p’s < 0.02) and *Cyfip1*^N/N^ mice derived from families with a paternal deletion showed a greater y-intercept than *Cyfip1*^N/N^ mice with a maternal deletion (p = 0.046). **(D,E):** In considering the effects of *Cyfip1* deletion and PO on a *Cyfip2*^J/J^ background, there was a main effect of Genotype [F(1,446) = 4.1; p = 0.04], Sex [F(1,446) = 11.8; p =0.0006], Day [F(1,446) = 21.3; p = 5.2 x 10^−6^] and importantly, there was a Genotype x PO interaction [F(1,454) = 10.4; p = 0.001] that reflected less PF consumption in *Cyfip1*^J/-^ mice with the maternal *Cyfip1*^J/-^ deletion (**D; \*** t(41) = 2.4; uncorrected p = 0.02 vs. *Cyfip1*^J/J^ on D11) but not paternal *Cyfip1*^J/-^ deletion (**E**). **(F):** We observed significant escalation in consumption in both *Cyfip1*^J/J^ wild-type groups [maternal: F(1,136) = 12.6; p = 0.0005; paternal: F(1,106) = 9.6; p = 0.003]. Neither *Cyfip1*^J/-^ group had a significant non-zero slope (both ps > 0.09). Moreover, *Cyfip1*^J/-^ mice with a paternal deletion had a greater y-intercept than all three of the other groups (**$;** all ps < 0.0004). **(G-L):** We also found a Sex x Genotype interaction [F(1,446) = 9.3, p = 0.002], a Gene x PO interaction [F(1,446) = 9.1; p = 0.003], and a Genotype x Sex x PO x Day interaction [F(1,446) = 5.7; p = 0.02]. To understand the source of these interactions, we next separated PO effects of *Cyfip1*^*J/-*^ by Sex. **(G-I):** In the females, we observed a main effect of Day [F(1,220) = 10.6; p = 0.001] and a Genotype x PO interaction [F(1,220) = 4.9; p = 0.03], **G,H)**. Moreover, both *Cyfip1*^*J/J*^ wild-type groups showed a significant escalation, regardless of PO (**I**; both ps < 0.02). Interestingly, although the *Cyfip1*^*J/-*^ mice with a paternal deletion had the greatest y-intercept (**$**; p < 0.0001), they had a negative slope of escalation (**I**; −3.4 x 10^−5^ ± 7.5 x 10^−5^) indicating initially higher PF consumption but a progressive decrease in consumption over time. **(J-L):** For males there was an effect of Genotype [F(1,226) = 13.6; p = 0.0003], Day [F(1,226) = 10.7; p = 0.001], a Genotype x PO interaction [F(1,226) = 5.7; p = 0.02], and a Genotype x PO x Day interaction [F(1,226) = 3.9; p = 0.049]. that was explained by maternally-deleted *Cyfip1*^*J/-*^ mice eating markedly less PF than their *Cyfip1*^J/J^ wild-type counterparts or, i.e., the induction of robust escalation in consumption in the wild-type *Cyfip1*^J/J^ males (**J, \***p < 0.03 vs *Cyfip1*^J/J^ on D16). In contrast, paternally-deleted *Cyfip1*^J/-^ mice showed no difference relative to their wild-type *Cyfip1*^J/J^ counterparts (**K**). In examining the slopes of escalation in all four groups, only the wild-type *Cyfip1*^*J/J*^ males coming from maternal deletion showed a significant escalation over time (**L**; p = 0.01). Data are presented as mean ± SEM.

**Figure 5.**
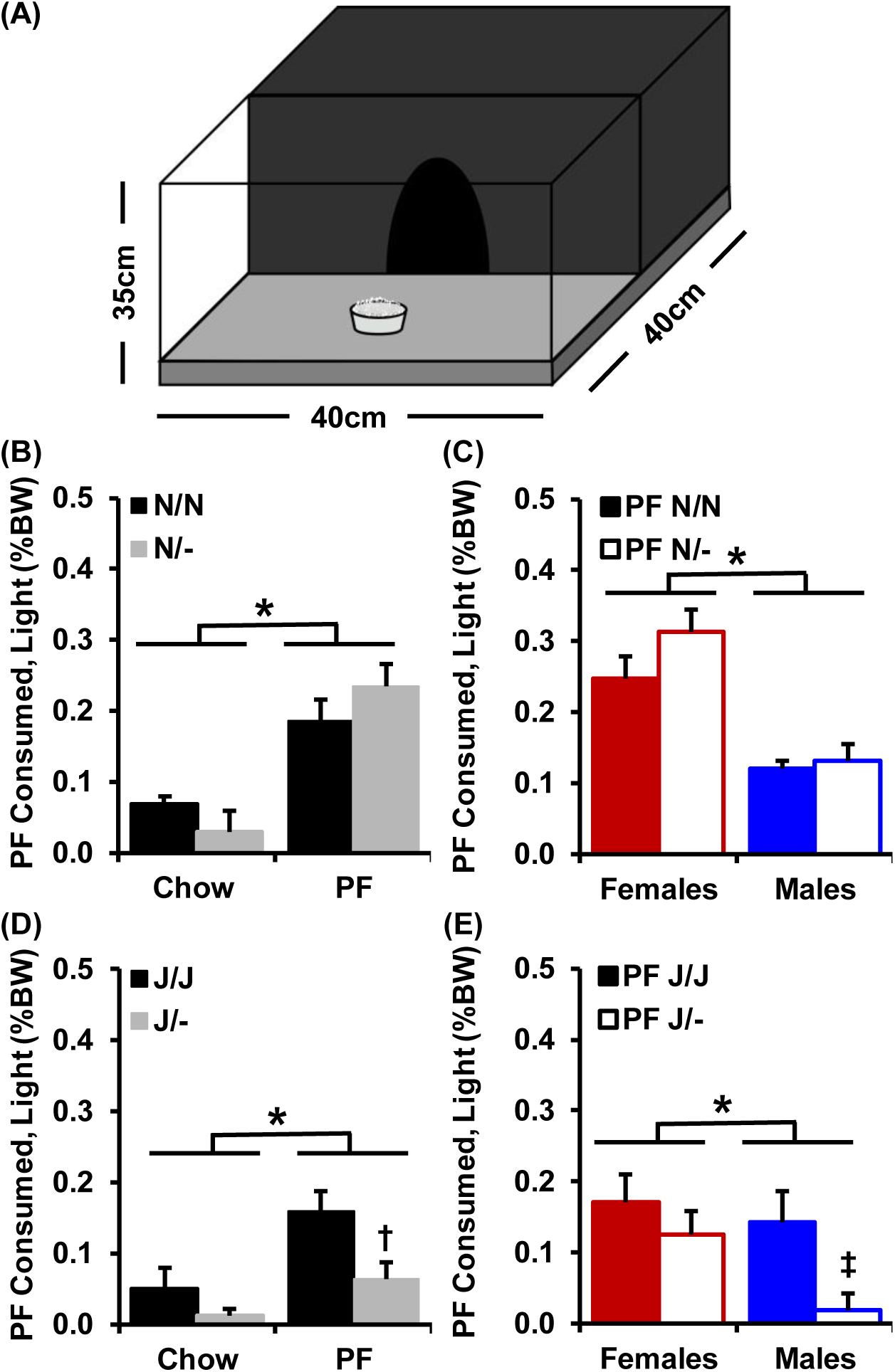
Compulsive-like PF intake in the light/dark conflict test in *Cyfip1*^N/-^ and *Cyfip1*^J/-^ mice. **(A):** A cartoon of the apparatus for the light/dark conflict test of compulsive-like PF consumption is shown. **(B):** For the *Cyfip2*^N/N^ background, there was a main effect of Training Treatment [**\***F(1,148) = 31.1; p = 1 x 10^−7^], indicating that PF-trained mice showed increased PF intake. However, there was no effect of Genotype [F(1,148) = 0.1; p = 0.7] or interaction of Genotype with Training Treatment on consumption [F(1,148) = 2.3; p = 0.13]. **(C):** In examining PF-trained mice alone, females showed increased intake [**\***F(1,72) = 13.6; p =0.0004]; however, there was no effect of Genotype [F(1,72) = 1.2; p = 0.3] or Genotype x Sex interaction [F(1,72) = 0.7; p = 0.4]. **(D):** For mice on the *Cyfip2*^J/J^ genetic background, there was a main effect of Training Treatment [*F(1,132) = 10.4; p = 0.002], indicating greater PF intake in PF-trained mice. There was also a main effect of Genotype [F(1,132) = 8.9; p = 0.003]. PF-trained *Cyfip1*^*J/-*^ mice consumed less than PF-trained *Cyfip1*^*J/J*^ mice [**†;** t(75) = 2.5; p = 0.02]. **(E):** In considering only PF-trained mice, females showed greater overall intake [**\***F(1,69) = 4.1; p =0.047] and *Cyfip1*^J/-^ mice showed overall less intake [F(1,69) = 4.9; p = 0.03]. However, only male *Cyfip1*^*J/-*^ mice consumed less than male *Cyfip1*^*J/J*^ mice [**‡**; t(37) = 2.7; p = 0.01]. Data are presented as mean ± SEM.

### Reduced transcription of *Cyfip1* but not *Cyfip2* or *Magel2* in the hypothalamus of *Cyfip1*^+/-^ mice

We hypothesized that the PO-and genetic background-dependent effects of *Cyfip1* deletion on PF intake could involve differences in hypothalamic gene transcription of *Cyfip1* and perhaps *Cyfip2*^8^. We also examined *Magel2* which is a nearby imprinted gene that is located within the syntenic, canonical (Type II) PWS locus and has been implicated in changes in eating behavior in mouse models of PWS^55^ and PWS-like hyperphagia in humans^54,56,57^. **Supplementary Table 3** lists the qPCR results as a function of both *Cyfip1* haploinsufficiency and PO. As expected, *Cyfip1* deletion significantly reduced gene transcription. When assessed on the *Cyfip2*^N/N^ background, the reduction in *Cyfip1* transcription in *Cyfip1*^*N*/-^ mice was similar following maternal versus paternal *Cyfip1* deletion (**Supplementary Table 3A**). However, when assessed on the *Cyfip2*^J/J^ background, the decrease in *Cyfip1* expression was enhanced following maternal *Cyfip1* deletion (**Supplementary Table 3B**) which was consistent with the enhanced reduction in PF consumption in maternally-deleted *Cyfip1*^J/-^ mice on the *Cyfip2*^J/J^ background (**Fig.4**). Finally, there was no effect of *Cyfip1* Genotype or PO on transcription of *Cyfip2* or *Magel2* **(Supplementary Table 3)**.

### Reduced CYFIP1 protein in *Cyfip1*^+/-^ mice depends on PO

We next investigated the effect of *Cyfip1* haploinsufficiency on protein expression in *Cyfip2*^J/J^ mice following completion of BE training. In the hypothalamus, we found a decrease in CYFIP1 protein in paternally-deleted *Cyfip1*^J/-^ mice **(Supplementary Fig.5A)**. When females and males were considered separately, we found a similar decrease in females resulting from paternal deletion **(Supplementary Fig.5B)**, but no difference in males **(Fig.5C)**. In contrast, in the nucleus accumbens, CYFIP1 expression was reduced in maternally-deleted *Cyfip1*^J/-^ mice **(Fig.5D)**. When we considered females and males separately, we did not detect any differences in CYFIP1 protein **(Fig.5E,F)**.

## DISCUSSION

*Cyfip1* haploinsufficiency increased OC-like behavior on two different *Cyfip2* genetic backgrounds (**Figs.1-2**) and altered PF consumption and *Cyfip1* gene expression, depending on genetic background (*Cyfip2*), Sex, and PO (**Figs.3-5**). These findings identify a significant contribution of reduced CYFIP1 expression to OC-like behaviors and PF intake that has relevance for neurodevelopmental disorders, including Type II PWS or the PWP (**FXS**). The selective increase in sweetened PF but not Chow intake during training (**Fig.3**) and during the test for compulsive-like eating (**Fig.5**) is consistent with increased preference for sweetened PF in PWS ^62-64^.

### OC-like behavior and PF intake in *Cyfip1*^+/-^ mice

The selective increase in OC-like but not anxiety-like behavior following *Cyfip1* deletion (**Fig.2**; **Supplementary Table 2**) is consistent with a lack of genetic correlation between marble burying and anxiety and supports marble burying as a repetitive, perseverative-like behavior^45^. Nevertheless, there is likely an anxiety-like component to marble burying^65^ as there is with OC behaviors in humans. For the *Cyfip1*^J/J^ genetic background, the increase in head-dipping behavior in the hole board task in *Cyfip1*^J/-^ mice further supports an increase in OC-like behaviors associated with *Cyfip1* haploinsufficiency^66^.

An increase in marble burying can predict BE^67,68^ and thus, our results suggest that human genetic polymorphisms affecting CYFIP1 expression could modulate OC behavior and risk for eating disorders^41,42^. OC behaviors are associated with both BE^37,38^ and PWS hyperphagia^40^. However, in our studies, there was no clear relationship between OC-like behaviors and PF intake. Although *Cyfip1* haploinsufficiency increased marble burying on both backgrounds (**Fig.2**) and increased PF consumption on the *Cyfip2*^N/N^ background, it *decreased* PF consumption on the *Cyfip2*^J/J^ background (**Fig.3**) while on the same background increasing OC-like head-dipping behavior (**Supplementary Fig.1**). Thus, our results show dissociable effects of *Cyfip1* deletion on OC-like behavior and the direction in modulation of PF intake. Notably, PWS patients show an increase in non-food OC behaviors, which is exacerbated in Type I PWS patients (who all have the *CYFIP1* deletion)^16,18,28-30^. *CYFIP1* haploinsufficiency could conceivably enhance OC behaviors and affect eating behaviors through separate, independent neural mechanisms. Additionally, although *CYFIP1* deletion is associated with the more severe, Type I form of PWS, we are not aware of any studies showing increased severity of hyperphagia in Type I PWS. Thus, CYFIP1 genotype could act more generally as a modifier of PWS hyperphagia rather than distinguishing PWS subtypes.

### Effect of *Cyfip1* deletion on PF intake depends on genetic background

As predicted, mice on a *Cyfip2*^N/N^ versus *Cyfip2*^J/J^ background showed greater overall PF consumption **(Fig.3B-E)**^69^. Additionally, *Cyfip1*^N/-^ mice showed a greater increase in PF consumption and escalation (**Fig.3D,E**). In contrast, *Cyfip1*^J/-^ mice on the lower PF-consuming *Cyfip2*^J/J^ background showed a *decrease* in escalation of PF intake (**Fig.3D,E**). This result was driven by the surprising induction of BE in wild-type *Cyfip1*^J/J^ males (**Fig.3F,G**), specifically males derived from maternal *Cyfip1*^J/-^ deletion (**Fig.4J,L**). The escalated intake in wild-type *Cyfip1*^J/J^ males was opposite to the low-level BE on the *Cyfip2*^J/J^ background^8^ and is inconsistent with a lack of BE in the parental B6J strain, especially males^8,35^. While we bred the *Cyfip2* locus to be fixed for the J allele, heterozygous N alleles segregating elsewhere in the genome could contribute to BE in *Cyfip2*^J/J^ mice. In support, F2 mice possessing the *Cyfip2*^J/J^ genotype showed greater PF consumption than the parental B6J strain^8^. However, heterozygosity at other loci cannot fully explain the current results, since only a specific subset of wild-type males (maternal *Cyfip1*^J/-^ - derived) showed an anomalous induction of escalation in consumption. Thus, wild-type *Cyfip1*^J/J^ males could be especially sensitive to social influences of maternal-pup and/or pup-pup interactions in the maternally-deleted *Cyfip1*^J/-^ environment, ultimately explaining escalation of PF intake.

In stark contrast to male *Cyfip1*^J/-^ mice with the *maternal* deletion, female *Cyfip1*^J/-^ mice (**Fig.3F,G**) with the *paternal Cyfip1*^J/-^ deletion showed a markedly enhanced, initial consumption of PF that obfuscated detection of escalation over time (**Fig.4H,I**). In considering sex differences in PWS, male PWS patients with 15q11-q13 deletions showed greater food-related preoccupation with food, impaired satiety, and other food-related negative behaviors in the absence of differences in OC or other non-food-related behaviors^70^. It is unclear whether sex differences in food-related problems differ depending on Type I versus Type II PWS deletions. In addition to behavioral differences, female PWS patients showed higher levels of circulating insulin and HOMA-IR, indicating greater insulin resistance and decreased adiponectin levels^71^. Together, our results illustrate the importance of considering Sex when investigating maternal versus paternal gene haploinsufficiency in eating behavior and the potential relevance to PWS in humans.

### Interactions of PO and offspring genotype in offspring behavior

The increase in PF consumption in all offspring from paternal *Cyfip1*^N/-^ deletion (**Fig. 4B vs. Fig.4A**) and the robust increase in PF consumption in wild-type *Cyfip1*^J/J^ males derived from maternal but not paternal *Cyfip1*^J/-^ deletion (**Fig.4J-L**) highlight potential genetic interactions with social environment in explaining behavioral variance^72^. Maternal versus paternal *Cyfip1* deletion could affect social interactions with the dam and sire or with the pups. Of direct relevance to *Cyfip1*, maternal deletion of the *Fmr1* gene (coding for FMRP) can induce neurobehavioral phenotypes in wild-type offspring and enhance phenotypes in mutant offspring, including locomotor hyperactivity, reduced behavioral response to D2 dopamine receptor activation, and enhanced behavioral response to GABA-B receptor activation^73^. In addition, males can demonstrate paternal pup retrieval^74^ and thus, *Cyfip1* deletion could also affect sire-pup contact. Given the association between *CYFIP1* deletion and social deficits in both 15q11.2 MDS and PWS but also FXS, autism, and schizophrenia^14^, it is plausible that *Cyfip1* deletion in the dam or sire affects social dynamics in the offspring in a PO-specific and genotype (offspring)-dependent manner, leading to long-term neurobehavioral effects.

### Hedonic hypothesis of *Cyfip1* modulation of PF intake

Despite differences in PF consumption as a function of *Cyfip1* Genotype, PO, Sex, and Genetic Background, there were no genotypic differences in body weight (**Supplementary Fig.2**), suggesting that *Cyfip1* haploinsufficiency could modulate PF intake through a non-homeostatic, perhaps hedonic mechanism involving altered sensory or affective processing of sweetened PF ^12,13^. The selective changes in PF consumption and conditioned reward (at least in *Cyfip1*^N/-^ mice; **Supplementary Fig.3A**) support the hedonic hypothesis and are consistent with enhanced preference for sweetened food in individuals with Type I PWS^62-64^. The *Cyfip2*^N/N^ mutation is associated with cocaine neurobehavioral sensitivity and plasticity^9^ and compulsive-like BE^8^. Furthermore, differences in in *Cyfip2* mRNA expression were genetically correlated with differences in cocaine self-administration^75^. Here, we observed PO-dependent decreases in CYFIP1 protein in both nucleus accumbens and hypothalamus (**Supplementary Fig.5**). Previous transcriptome analysis of the striatum from *Cyfip2*^N/-^ versus *Cyfip2*^N/N^ genotypes identified “morphine addiction” and “cocaine addiction” as two of the top five KEGG enrichment terms^8^. FMRP, a major interacting protein with CYFIP1/2, is involved in reward processing^76^ and cocaine neurobehavioral plasticity^77^. We hypothesize that *Cyfip1* deletion/polymorphisms could alter the hedonic effects of PF consumption via the dopaminergic mesolimbic reward circuitry and interact with other haploinsufficient genes underlying hypothalamic, homeostatic mechanisms of hyperphagia in PWS to modulate food intake^78,79^.

### Sex-dependent PO effects on Cyfip1 expression and behavior

There is no published evidence that *Cyfip1* is imprinted and while our analysis of *Cyfip1* transcript and protein levels indicate PO-dependent effects, the direction was not always consistent with maternal imprinting and was dependent on Sex and the particular brain region (**Supplementary Table 3; Supplementary Fig.5**). A recent study of nearly 100 phenotypes showed that most complex traits exhibit PO effects and that non-imprinted KO alleles (e.g., *Cyfip1*) can induce extensive PO effects by interacting in *trans* with imprinted loci throughout the genome to affect gene networks^80^. If a *trans*-acting genomic mechanism underlies the effects of *Cyfip1* haploinsufficiency, it would have to be co-inherited faithfully with the maternal or paternal deletion to explain the PO effects on behavior. An obvious candidate mechanism could involve inheritance of sex-dependent gene expression originating from sex chromosomes (and potential sex chromosome variants between substrains that would explain genetic background-dependent effects) that interacts with *Cyfip1* deletion. This is particularly relevant to CYFIP proteins given that *Fmr1*, which codes for FMRP, the interacting protein of CYFIP that regulates protein translation and is located on the X chromosome. The human *FMR1* gene undergoes X-inactivation^81^; thus, females and males should have equivalent levels of gene dosage, transcription, and translation of FMRP. Nevertheless, 10-25% of human genes and 3-7% of mouse genes show variable degrees of X inactivation^82,83^. *Fmr1* could undergo variable x-inactivation depending on the tissue and time point. Furthermore, genetic polymorphisms between B6J and B6NJ on the X chromosome could affect the expression of genes that act as modifiers of *Cyfip1* transcription or modifiers of X-inactivation and account for the background-dependent PO effects of *Cyfip1* deletion on behavior. The use of the four core genotypes model (XX, XY, XX-male, XY-female) could be used to test the involvement of sex chromosomes in PO-and Sex-dependent effects of *Cyfip1* deletion on PF intake - a recent study using this genetic model identified a contribution of sex chromosomes to operant reinforcement for PF^84^.

## Conclusion

Our preclinical findings provide evidence that reduced CYFIP1 expression could contribute to OC and eating behaviors in PWS and other neurodevelopmental disorders. Future genomic studies of multiple brain regions, cell types, and developmental time points could inform molecular mechanisms of eating behaviors on different genetic backgrounds and potential interaction of *Cyfip1* deletion with gene expression on sex chromosomes. Furthermore, PO effects of *Cyfip1* deletion could be tested for behavioral and genomic interactions with imprinted genes in PWS models of hyperphagia (e.g., deletion of *Magel2* or *Snord116*). Such efforts could improve upon existing PWS models that, like our findings, have historically lacked obesity^85^ and could inform pharmacotherapeutic treatment of eating behavior tailored to a particular subtype of neurodevelopmental syndrome.

## ACKNOWLEDGEMENTS

Funding was provided by NIH/NIDA R21DA038738 (C.D.B.), NIH/NIDA R01DA039168, NIDA Diversity Scholars Network (F.R.), Burroughs-Wellcome Fund Transformative Training Program in Addiction Science (TTPAS; 1011479), NIH/NIGMS T32GM008541, and NIH/NIDA U01DA041668 (V.K.). We thank Dr. Rachel Wevrick for providing us with the primer sequences used for gene expression analysis of *Magel2*. We would also like to acknowledge Dr. Lynn Deng and Matthew Au of the Boston University Analytical Instrumentation Core Facility (S10OD023663) for their support in conducting the qPCR studies.

